# Direct link between convergent evolution at sequence level and phenotypic level of septal pore cap in Agaricomycotina

**DOI:** 10.1101/2023.03.02.530715

**Authors:** Tomoyo Iizuka, Masafumi Nozawa, Kazuho Ikeo

## Abstract

Several apparently homologous morphological characters are known to have independently evolved in different lineages multiple times. However, the genetic backgrounds of such morphological convergences are not well understood. To detect any correlated genes potentially responsible for morphological convergence at the phenotypic level, we focused on the morphology of the septal pore cap (SPC), which is involved in mycelia’s complex multicellularity in fungi. SPCs are classified into three morphological types: perforate, imperforate, and vesiculate. To understand what evolutionary events occurred at the sequence level during morphological convergence of perforate SPCs in Agaricomycotina, we examined sequence differences between species with different SPC types by comparative genomic analysis. If sequences from species with perforate SPCs formed a cluster, the associated gene might be involved in generating perforate SPCs in multiple lineages. Based on this assumption, we detected eight candidate genes, including an SPC-related gene, *spc33*. The results showed that some amino acid substitutions independently occurred in both lineages in which species with perforate SPCs emerged. From these results, we speculate that the amino acid substitutions in *spc33* were critical in the emergence of perforate SPCs in multiple lineages. We also found that *spc33* evolved just before imperforate SPC emergence based on homology search of *spc33* and the species phylogeny. These findings illustrate the first step to clarifying the genetic basis of SPC morphological evolution. Our study contributes to both clarifying the genetic basis of morphological convergence and pioneering our understanding of fungal evolutionary morphology.

## 1. Introduction

An important problem in evolutionary biology is determining what changes at the genome sequence level have produced morphological diversity. The study of convergent evolution, i.e., the independent evolution of similar phenotypic changes in different lineages, is thought to provide insight into hotspots of key mutations for phenotypic differentiation (Stern and Orgogozo, 2009). In addition, focusing on morphological convergence provides clues for understanding the relationship between phenotypic evolution affected by natural selection and sequence evolution that can be described by the protein adaptive landscape (Castoe et al., 2010; Stern 2013; Almén et al., 2016). Therefore, understanding convergent evolution at the sequence level is also very important for clarifying which substitutions produced convergent evolution at the phenotypic level. However, genome-wide surveys of convergent evolution are relatively rare and more empirical genome-scale studies are needed to elucidate the genetic basis of morphological convergence.

One such example of morphologically convergent traits is fungal septal pore caps (SPCs), which are involved in the mycelia’s complex cell-to-cell communication (Jedd, 2011; Nguyen et al., 2017). It was suggested that SPCs were newly derived from the endoplasmic reticulum (ER) because SPCs were stained by some ER-targeting markers (van Driel et al., 2008; Müller et al., 1999) and the base of SPCs are connected to the ER (Moore, 1975; Müller et al., 1998a, 2000). Thus, understanding the genetic background of SPC provides fundamental insight into the origin of multicellularization in fungi and the mechanism of organelle specialization of SPC.

We investigated three morphological types: vesiculate, imperforate, and perforate SPC. In Agaricomycotina, taxa of the most ancestral lineage have vesiculate SPCs, whereas taxa of all other lineages have either imperforate or perforate SPCs. A previous phylogenetic study showed that perforate SPCs independently evolved from imperforate SPCs multiple times (van Driel et al., 2009); therefore, perforate SPCs evolved under morphological convergence. The different SPC types result in different cytoplasmic transport functions inside the mycelium (Bracker and Butler, 1964; Moore and Marchant, 1972), which indicates that SPC types evolved through adaptation to transport various substances. These SPC types are highly conserved at the order level (Hibbett, 2006; van Driel et al., 2009; Oberwinkler et al., 2013; Hibbett et al., 2014) and understanding the genetic background of SPC morphological differentiation can substantially contribute to understanding the evolution of higher taxa in Basidiomycota.

Previous SPC studies reported on their functions, fine features, and applicability to fungal classification (Bracker and Butler, 1964; Moore and Marchant, 1972; Lisker et al., 1975; Hibbett, 2006; van Driel et al., 2009); however, despite the long history of such morphological studies on SPCs, the genetic basis remains unclear. Previously, most studies focused on visible phenotypes when reporting substitutions associated with morphological convergence by whole-genome analysis. For example, Hu et al. (2017) found a candidate gene for the evolution of a pseudo-thumb in pandas using positive selection as an indicator. Colosimo et al. (2005) found that the Eda pathway is involved in the phenotypic convergence of armor plate in sticklebacks. Therefore, the genomic background associated with morphological convergence of SPC, that is hard to tackle using reverse or forward genetics approaches, could also be elucidated using genomic approaches.

In this study, we searched signatures of sequence evolution that had evolved with morphological convergence of SPC by comparative genomics. We collected publicly available whole-genome sequences from the Joint Genome Institute portal (JGI, http://www.genome.jgi.doe.gov). Because of their importance for taxonomic classification (Wells, 1994; Müller et al., 1998b, 2000; Lutzoni et al., 2004; Hibbett et al., 2014), data on morphological types of SPC have accumulated. Although the available set of genomic and morphological data were limited, we successfully inferred evolutionary events that corresponded to morphological convergence at the amino acid sequence level using datasets of representative 12 Agaricomycotina species. Here we report that convergent evolution had occurred in sequence level as well as phenotypic level. In particular, amino acid substitutions in the SPC-related gene *spc33* accumulated and directly correlated with the morphological difference between imperforate and perforate SPCs. Therefore, the gene that plays a role in SPC formation may also be involved in the evolution of SPC morphology. These findings provide the first step to clarify the genetic basis of the morphological evolution of SPC and possible contribution of limited combination of key substitutions to the morphological convergence.

## 2. Materials and Methods

### 2.1 Genome sequences

The coding sequence data for 12 species of Basidiomycota, which included 11 representatives of Agaricomycetes (six perforate species and five imperforate species) and one representative of Dacrymycetes (imperforate type), were used for this study (Table 1). These species are representative of major Basidiomycota species. All sequence data were obtained from the JGI of the United States Department of Energy.

**Table 1.**
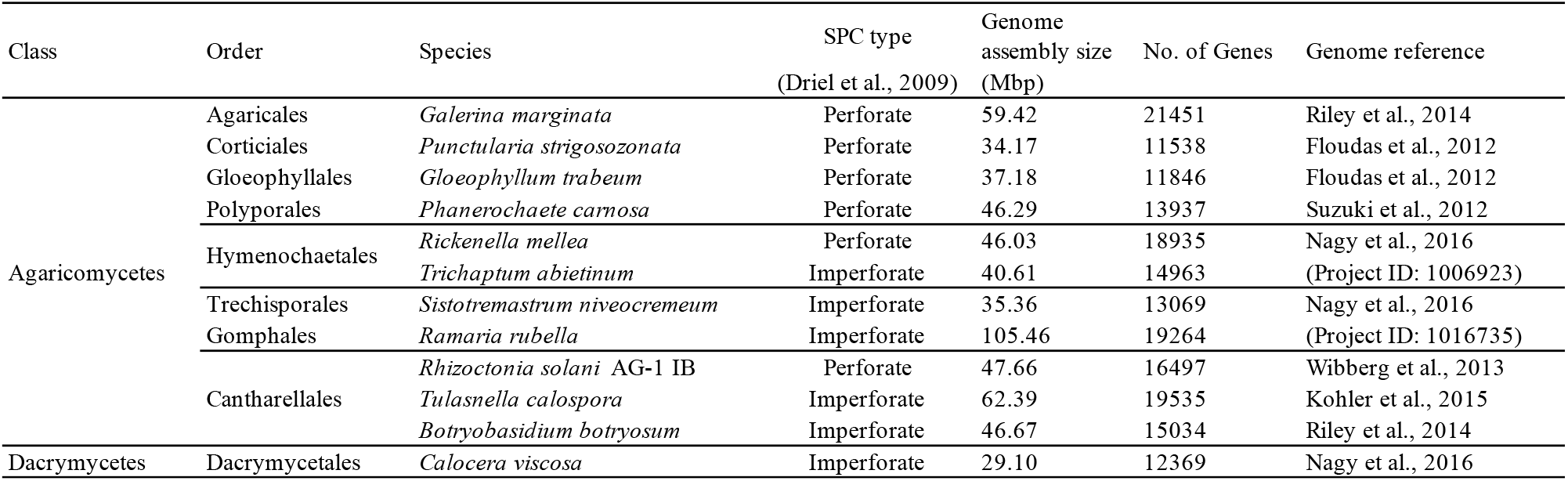
Taxonomic classification and data sources for species included in this study.

### 2.2 Ortholog identification

To identify orthologs, reciprocal best hits (RBH) analysis (Tatusov et al., 1997; Bork et al., 1998; Overbeek et al., 1999) was conducted. We performed all-vs-all comparison with BLAST software (blastp, NCBI-blast-2.2.30+) using the coding sequences that remained after discarding sequences with internal stop codons. The RBH were extracted using original Perl scripts. These scripts are available on a website of Laboratory for DNA Data Analysis of National Institute of Genetics (http://molevo.sakura.ne.jp/DnaData_lab/).

### 2.3 Detection of convergent substitutions

We conducted multiple alignments of candidate genes and extracted key sites for morphological convergence at a single amino acid site level from the genomes of 12 Basidiomycota species. The Multiple alignment was performed using MUSCLE v3.8.31 (Edger, 2004). To examine the possibility that the same evolutionary event occurred at both phenotypic and sequence levels, we searched convergent substitutions from the amino acid sequence of the orthologous gene dataset by conducting PCOC analysis (Rey et al., 2018) and a traditional method referred from Zhang and Kumar (1997). In our traditional method, the ancestral sequences of all orthologs were also estimated using the ML approach implemented in MEGA6-CC (Tamura et al., 2013). A substitution was regarded as a convergent substitution if an amino acid independently changed to the same amino acid in the lineage to perforate species. This “number of convergent substitutions” was calculated for each ortholog. Then, we calculated the number of convergent substitutions per whole substitutions for each ortholog; for example, when an ortholog carries 3 convergent substitutions and 100 whole substitutions, it becomes 3 ÷ 100 = 0.03.

### 2.4 Phylogenetic analysis of orthologous gene dataset

Candidate genes associated with the morphological differences between imperforate and perforate SPCs were extracted from orthologous genes by conducting phylogenetic analyses and assessing their topologies. This approach is based on a working hypothesis that the same genes repeatedly contribute to the emergence of perforate SPCs independently in multiple times. Thus, we extracted genes that showed correlated changes with morphological traits by constructing a gene tree for each ortholog.

The same multiple alignments as described in 2.3 were used in this analysis. Phylogenetic gene trees for each ortholog were generated using the neighbor-joining (NJ) (Saitou and Nei, 1987) and maximum-likelihood (ML) methods (Felsenstein 1981) in MEGA6-CC (Tamura et al., 2013). We then checked the topology of the trees. A concatenated tree using a single copy orthologous gene dataset was also constructed using the NJ method to confirm the phylogenetic relationships of the 12 fungal species. We also confirmed reliability of our gene tree by conducting the approximately unbiased (AU) test (Hasegawa, 2002) with IQ-TREE v1.16.12 (Chernomor et al., 2016). In the topological analysis, we searched for gene trees that all the genes from species with perforate SPCs were in the same clade and all those with imperforate SPCs were in the other clade. When the topologies of both the NJ and ML trees differed from the species phylogeny, the gene was considered as a candidate gene (Fig. 1). The obtained genes were annotated using BLAST (Johnson et al., 2008) against five databases, NCBI non-redundant protein sequences, UniProt (259, 2017-04) (The UniProt Consortium, 2017), GeneOntology (2017-04-11) (The Gene Ontology Consortium, 2000), *Saccharomyces* Genome Database (R64.2.1, 2014-11-18) (Cherry et al., 2012), and PomBase (30_62, 2017-01-30) (Wood et al., 2012).

**Fig. 1.**
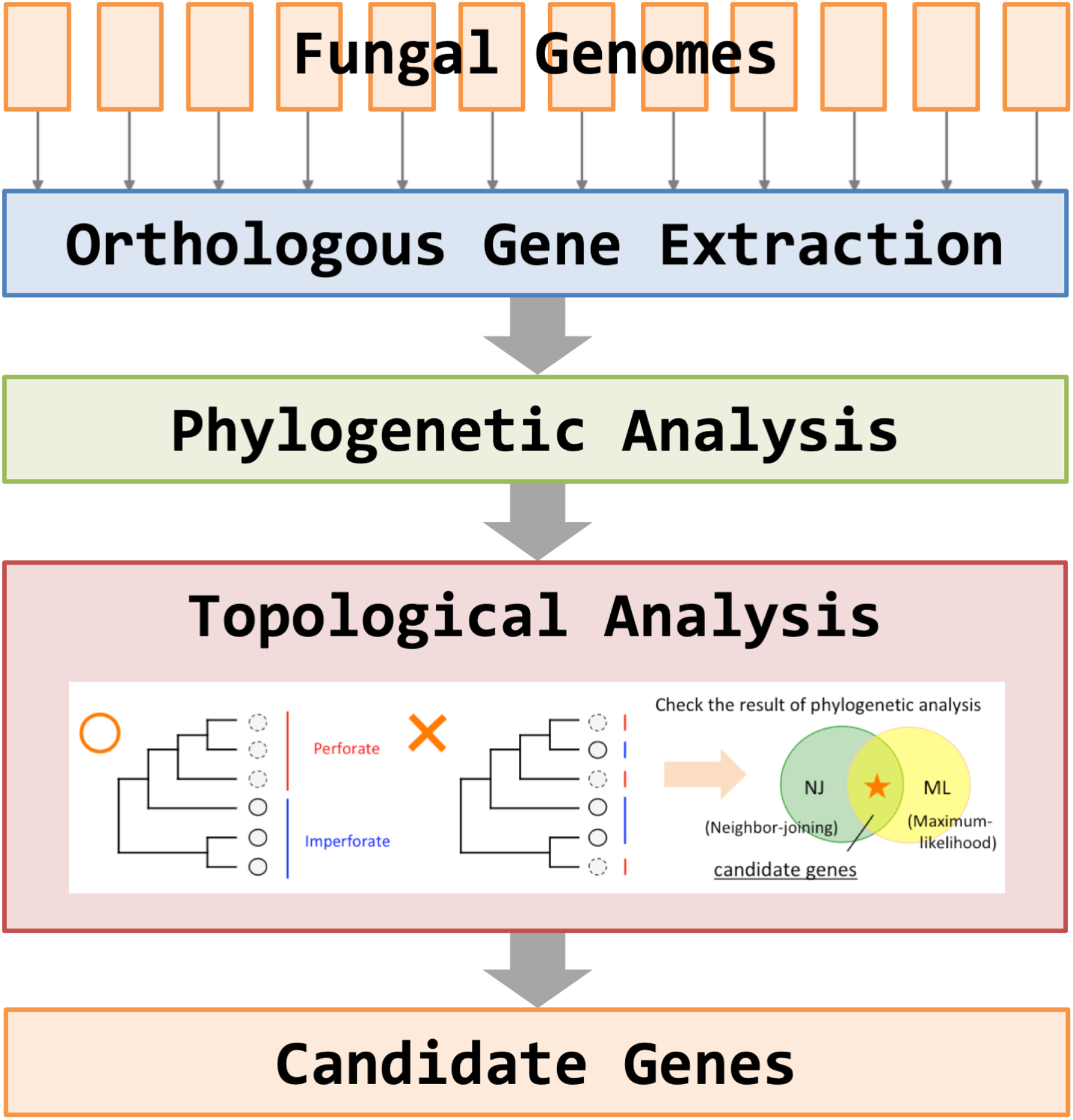
Candidate gene extraction algorithm.

## 3. Results

### 3.1 Convergent amino acid evolution detected from the orthologous gene dataset

The RBH analysis yielded 2560 single-copy orthologs. From 2560 genes, 111,654 and 2020 convergent sites were detected by PCOC analysis and our traditional method, respectively. (Supplementary Fig. 1). 84.6% of the sites detected by the traditional method were also supported by the PCOC method. For genes with convergent substitutions, the number of convergent sites detected by PCOC method increased in proportion to the total number of substitutions in the ortholog (R^2^ = 0.9752) (Supplementary Fig. 2).

### 3.2 Phylogenetic gene trees and candidate gene extraction

Our concatenated tree using these orthologs was consistent with the current species phylogeny (Fig. 2a). Gene trees constructed with the NJ method (Fig. 2b) and ML method (Fig. 2c) showed that eight genes of the perforate species *Rhizoctonia solani* AG1-IB (Rhiso; Wibberg et al., 2013) clustered with those of other perforate species from another lineage (Fig. 2, Table 2, Supplementary Fig. 3). Seven of these genes were annotated as known genes and another gene as unknown based on the referred databases.

**Table 2.** List of eight candidate genes.

|  | BS NJ <sup>1)</sup> | BS ML <sup>1)</sup> | model <sup>2)</sup> | aa <sup>3)</sup> | Gene symbol | Full gene name |
| --- | --- | --- | --- | --- | --- | --- |
| #1 | 40 | 22 | LG+G | 596 | *spc33* | 33kDa protein purified from SPCs |
| #2 | 11 | 10 | LG+G | 468 | DSD1 | D-serine dehydratase |
| #3 | 40 | 46 | rtREV+G | 103 | SDH8 | Succinate dehydrogenase |
| #4 | 15 | 12 | LG+G | 318 | n/a | Uncharacterized protein |
| #5 | 9 | 10 | LG+G | 98 | SUMO | Small ubiquitin-like Modifier protein |
| #6 | 37 | 38 | JTT+G+I | 454 | ELP4 | Elongator Acetyltransferase Complex Subunit 4 |
| #7 | 9 | 12 | LG+G+I | 271 | DRE2 | Fe-S cluster assembly protein |
| #8 | 21 | 12 | LG+G | 221 | DUS1 | tRNA-dihydrouridine synthase 1 |

1) Bootstrap (BS) value of the neighbor-joining (NJ)/maximum-likelihood (ML) tree supporting a single perforate SPC cluster.
2) Substitution model of the ML tree.
3) Amino acid length (*Rhizoctonia solani*).

**Fig. 2.**
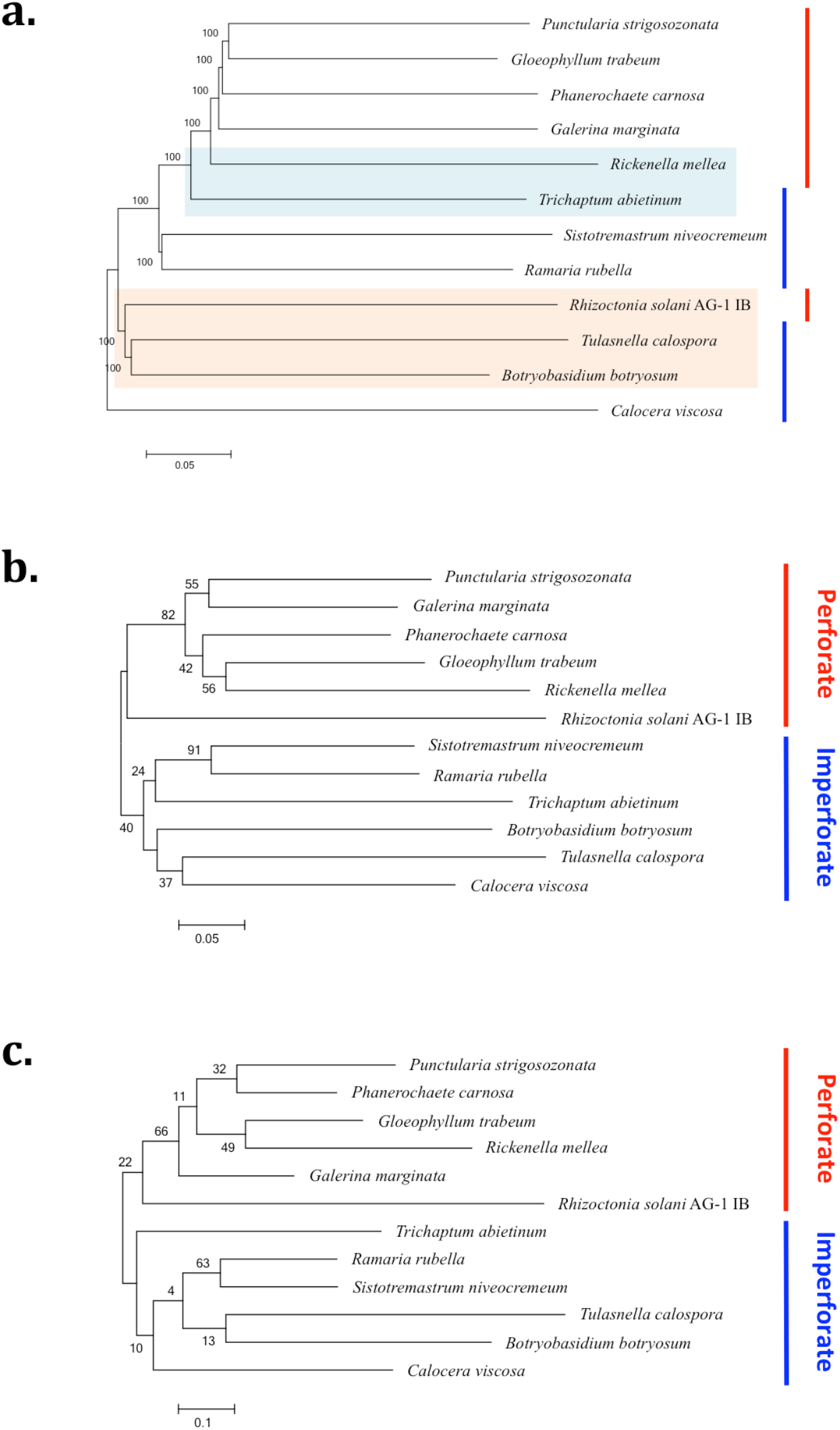
Phylogenetic relationship of 12 fungal genomes and correlated genes. (a) Phylogenetic tree based on the neighbor-joining method of concatenated sequences of 2560 orthologs. Numbers near internodes correspond to values of 1000 bootstrap replicates. *Hymenochaetales* (red) and *Cantharellales* (blue) were used for multiple species. (b, c) Gene trees of the correlated gene #1 based on (b) the neighbor-joining and (c) the maximum-likelihood methods. Phylogenetic gene trees of correlated genes #2–#8 were compiled in Supplementary Fig. 1. Scale bar at the bottom of each tree indicates the branch length.

### 3.3 The most reliable candidate gene *spc33*

One of the genes shown at 3.2 was identified as an SPC-related gene, *spc33* (Peer et al., 2007). The result of the AU test of *spc33* was *P* = 0.499, whereas that of our concatenated species tree was *P* = 0.024 with the appropriate amino-acid substitution model for the *spc33* gene tree (Supplementary Table 1). In addition, we found that convergent substitutions were accumulated in *spc33*. To test whether this observation was potentially random, we checked the number of convergent substitutions in all genes of Rhiso and other perforate lineages. The results showed that the proportion of convergent substitutions for spc33 was 0.3 × 10^−3^, which was top 1.3 percentile among all genes analyzed (Fig. 3). Unlike *spc33*, not all of the other 7 genes carried many parallel substitutions. These findings indicate that *spc33* has a high convergent substitution rate than other orthologs.

**Fig. 3.**
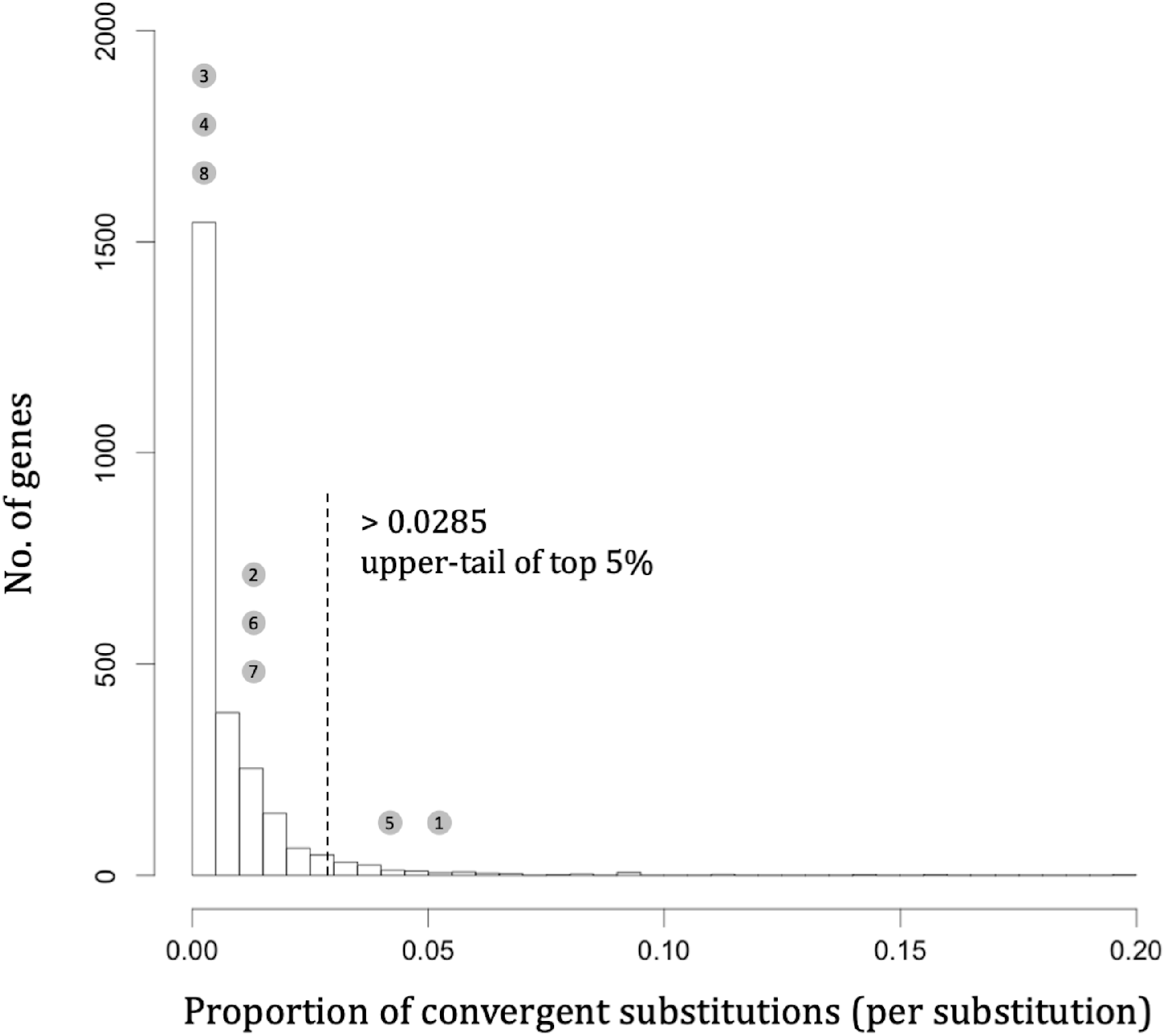
Frequency distribution of convergent substitution rates in 2560 genes. The x axis shows the proportion of convergent substitutions as described at 2.3. This showed that many genes do not have convergent substitution. Numbers 1 through 8 corresponds to those in Table 2.

### 3.4 Convergent evolution of *spc33* in amino acid and nucleotide sequence levels

We expected that if convergent evolution had also occurred at the amino acid sequence level, the phenotypic characterization of perforate SPCs may be regulated under a limited combination of molecular pathways. To investigate that, we searched for amino acid convergent substitutions in ancestral states of *spc33* from imperforate to perforate species. The results showed the same pattern of convergent amino acid substitutions at D354E, K357R, V359I, and P410R during evolution from perforate–imperforate branching point 1 (PI1) to the lineage including five perforate species (Perf1) and from perforate–imperforate branching point 2 (PI2) to Rhiso (Rhizoctonia solani AG-1 IB) (Fig. 4). Interestingly, three of the four sites were localized within the same short region of 17 amino acids, even though SPC33 is more than 300 amino acids in length. This finding indicates that sequence differences in this region of spc33 correspond to the morphological convergence of perforate SPCs. At the substitution K357R of *spc33*, both ancestral branching points PI1 and PI2 showed the same amino acid residue, lysine (Fig. 4a); alternatively, the estimated ancestral residue of site 357 was arginine at the ancestral branching point Perf1 (Fig. 4a) and the residue of Rhiso also changed to arginine at the emergence of this species (Fig. 4b). Also, this substitution pattern K357R from imperforate to perforate species was completely conserved in all fungal species (Fig. 4a). Therefore, the same amino acid substitution from lysine to arginine independently occurred when perforate species emerged. In other words, K357R and M/V359I still showed differences in the extant species (Fig. 4b, Supplementary Fig. 4); therefore, convergent evolution also occurred at the sequence level. Substitutions at D354E and P410R were also detected; however, these substitution patterns were not conserved in the extant species.

**Fig. 4.**
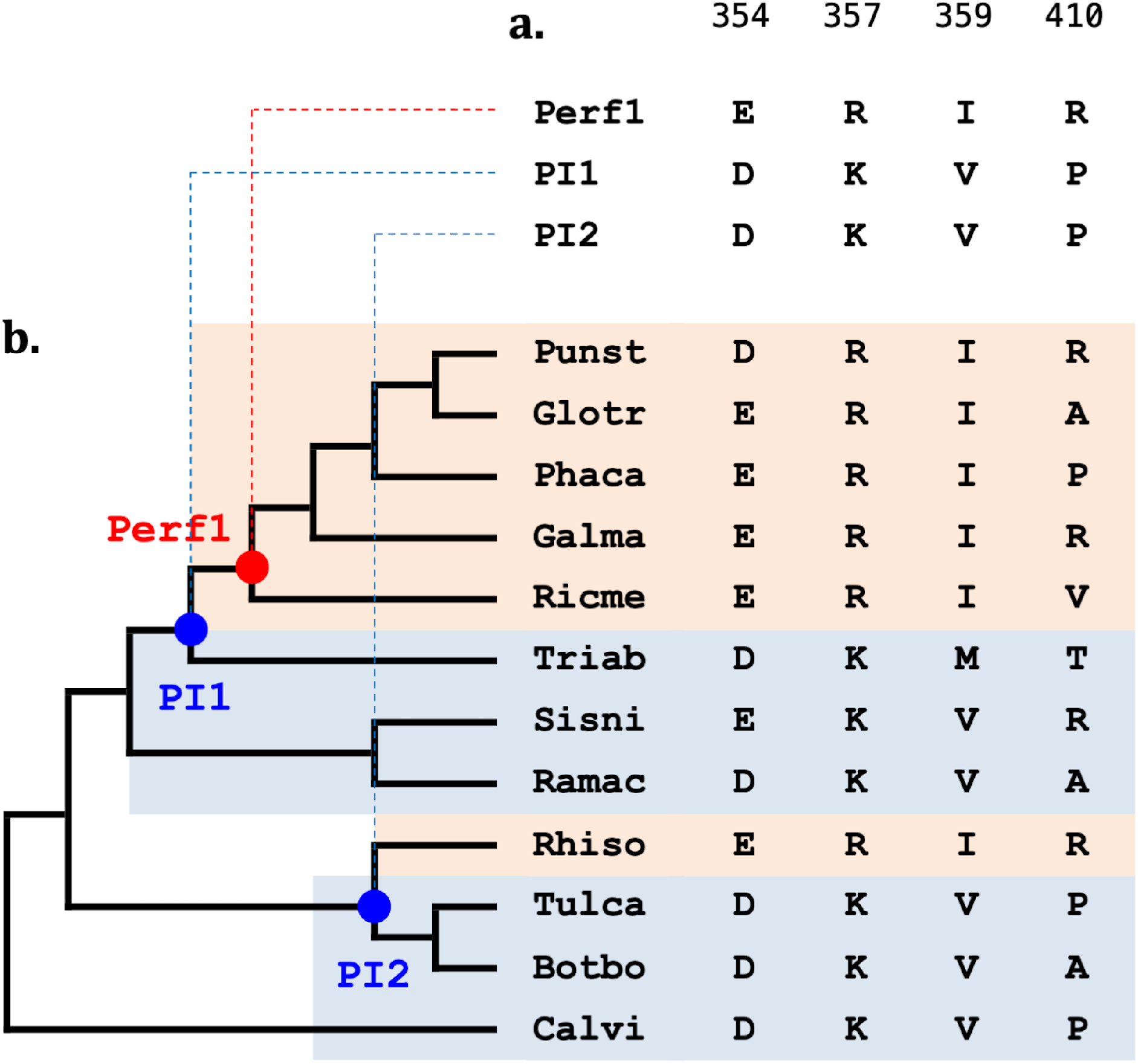
SPC type-specific convergent substitutions of *spc33* in perforate type species (red background) and imperforate type species (blue background). (a) Ancestral sequences of focal sites. The abbreviations of the branch names are as follows: Perf1, lineage including five perforate species; PI1, perforate–imperforate branching node 1; PI2, perforate–imperforate branching node 2. (b) SPC type-specific amino acid residues in extant species. The abbreviations of the species names are as follows: Punst, *Punctularia strigosozonata*; Glotr, *Gloeophyllum trabeum*; Phaca, *Phanerochaete carnosa*; Galma, *Galerina marginata*; Ricme, *Rickenella mellea*; Triab, *Trichaptum abietinum*; Sisni, *Sistotremastrum niveocremeum*; Ramac, *Ramaria rubella*; Rhiso, *Rhizoctonia solani*; Tulca, *Tulasnella calospora*; Botbo, *Botryobasidium botryosum*; Calvi, *Calocera viscosa*.

To understand substitution patterns in nucleotide sequences that resulted in amino acid substitution, we also checked site 357 at the nucleotide sequence level of both perforate lineages. Rhiso acquired the codon CGT (arginine) at site 357 (Fig. 5), and the two closest imperforate species (Tulca; *Tulasnella calospora* and Botbo; *Botryobasidium botryosum*) and the most basal species (Calvi; *Calocera viscosa*) use AAG (lysine). Other perforate species also use CG at the 1st and 2nd codons. This result indicates that the substitution pattern of all perforate species has changed to CGN (arginine) from AAR (lysine). Thus, even at the nucleotide sequence level, we observed the same substitution pattern from AAG to CGT (Fig. 5).

**Fig. 5.**
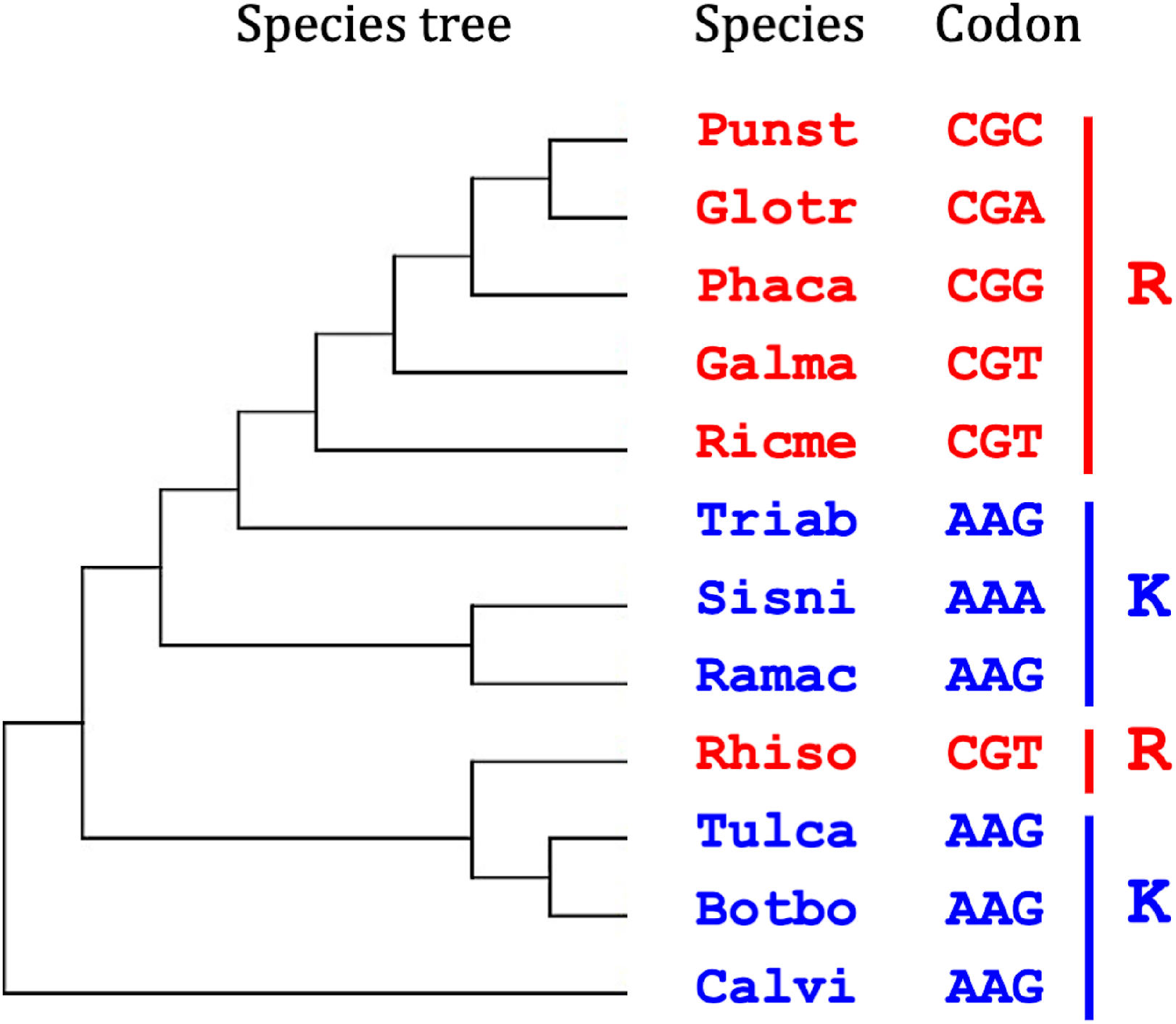
Nucleotide sequences of SPC type-specific site 357 in *spc33*.

### 3.5 Feature of *spc33* in taxa other than perforate and imperforate type species

To understand *spc33* sequences in other organisms, we searched for *spc33* homologs against whole sequence data available in the NCBI database and 41 fungal genome sequences in JGI, including the 12 fungal genomes tested in this study and genomes of species with vesiculate SPCs. Surprisingly, the blastp results with an E-value cutoff of 1E-5 showed that *spc33* was not found in any vesiculate species, whereas *spc33* was found in all imperforate and perforate species (Table 3). Moreover, *spc33* homologs were not found in organisms other than perforate and imperforate species (including other fungi, other eukaryotes, and prokaryotes). These presence/absence patterns of *spc33* indicate that *spc33* emerged before the divergence of imperforate SPCs from vesiculate SPCs. Furthermore, no known domain was found in *spc33* based on a Pfam 32.0 search.

**Table 3.** Presence/absence pattern of *spc33*. Numbers in parentheses indicate the number of fungal species tested.

|  | Vesiculate (2) | Imperforate (11) | Perforate (28) |
| --- | --- | --- | --- |
| Presence (%) | 0 | 100 | 100 |

## 4. Discussion

Van Driel et al. (2009) provided evidence for morphological convergence of SPCs based on species phylogeny. Here, we provide evidence for sequence convergence accompanying the morphological convergence of perforate SPCs. It was proposed that when there is convergent evolution at the phenotypic level, different genes or different sites of the same gene would lead to the same phenotype (e.g., Hoekstra and Nachman, 2003; Foote et al., 2015; Zou and Zhang, 2015; Corbett-Detig et al., 2020). In other words, there are multiple roadmaps that results in convergent phenotypic evolution. On the other hands, many genome-wide surveys reported concentration of convergent amino acid substitutions at the same sites in the same genes that potentially contribute to phenotypic convergent evolution (e.g., Nei et al., 1997; Shi and Yokoyama 2003; Natarajan et al. 2016; Hague et al., 2017; Hu et al. 2017). This means a few convergent substitutions can cause phenotypic convergence. Our findings support the latter scenario for the convergent evolution of perforate SPCs. Below, we discuss the indications for convergent evolution in fungal genome sequences and suggest the possible role of detected genes (especially *spc33*, the gene identified as SPC-related genes by van Peer et al. 2011) in the morphological convergence of SPCs.

Our analysis detected multiple amino acid sites that had occurred convergent evolution accompanied by morphological convergence of SPC, as well as key sites and clue genes. The PCOC approach has been successful in detecting convergent evolution of sequence level in PEPC protein among plant samples (Rey et al, 2018). However, in our dataset, the number of sites detected by PCOC analysis tended to be unnaturally high (Supplementary Fig. 1). In addition, for each gene, the number of convergent sites was proportional to the total number of substitutions in each gene, suggesting that PCOC analysis contains false positives (Supplementary Fig. 2). Therefore, it was necessary to conduct convergence site extraction with other criteria to detect the sites that might have more functional significance. Our approach has traditionally been used to select for sites strongly suggestive of functional variation and detects convergent sites using stricter criteria than PCOC (Zhang and Kumar, 1997). Here, we examine the entire gene sequence using these two approaches and found that 84.6% of the sites detected by the traditional method also detected by PCOC method, which uses less stringent criteria for determining convergence (Supplementary Fig. 1). These results indicate that our analysis was able to accurately detect convergent substitution sites.

Phylogenetic and topological analysis against orthologous gene dataset detected eight genes as candidates. Only these gene trees were clustered by SPC type (Fig. 2b, 2c). Honestly, bootstrap values of each phylogenetic gene tree of candidate genes are low. This is because we are assuming convergent evolution, means irrelevant sites that has not evolved accompanied by speciation must be contained the sequence data of candidate genes, and thus we cannot expect a high bootstrap value supporting convergent evolution. Furthermore, the results of AU test did not reject the topology of this tree (Supplementary Table 1, P = 0.499). This supported our assumption that the topology of the gene trees should be correlated with morphological traits and differed from the species phylogeny.

SPCs are known to interact with many intracellular substances, such as the ER, cytoplasm, and cytoskeleton (Bracker and Butler, 1964; Moore and Marchant 1972). Although we performed functional characterization of the candidate genes, we could not find an association with SPC in seven of eight genes. Therefore, the relationship between SPCs and these seven genes will be revealed when the relationship between these intracellular substances and SPCs is revealed. We also found that the two candidate genes were in the top 5% in terms of accumulation of convergent substitution sites (Fig. 3). One of them codes SUMO protein (Table 2). Due to the wide-ranging of SUMO’s functions (Gupta et al., 2020), further analysis is definitely needed to narrow down the possible involvement of SUMO in morphological changes of SPC at this point. Interestingly, there was one gene that had more convergent substitutions than the other genes and was also a candidate gene in the phylogenetic analysis, SPC-related gene *spc33*. It is one of three known SPC-related genes, *spc14, spc18*, and *spc33* (Peer et al.2010, van Driel et al., 2018). van Peer et al. (2010) demonstrated that *spc33* is a transmembrane protein of SPCs by examining *Schizophillum commune* (perforate type). When van Peer et al. constructed the *spc33*-knockout mutant of *S. commune*, the SPCs disappeared. It is very difficult to conclude that all pieces of evidence are entirely coincidental. By combining findings of their report with our findings, we successfully detected *spc33* as a gene correlated with the morphological convergence of SPCs. Furthermore, considering that BLAST analysis did not detect *spc33* from taxa other than perforate and imperforate species (Table 3), we propose *spc33* emerged just before the emergence of imperforate SPCs. This result supports Padamsee et al. (2012), who suggested that different genes other than *spc33* are involved in the SPC development in vesiculate species such as *Wallemia sebi*. Therefore, at this point *spc33* is the promising candidate gene for morphological evolution of SPCs, and it is biologically valid to discuss the convergent evolution of SPC morphology using *spc33* as a starting point.

We found that the specific SPC-associated convergent sites were located within amino acid sites 354-359, a short region of *spc33* (Fig. 4). This phenomenon led us to further question whether these substitutions have a functional role for morphological differentiation of SPCs. It is known that *spc33* is responsible for constructing SPCs (Peer et al., 2010). Additionally, we determined that *spc33* evolved to the same pattern of amino acid sequences during perforate SPC evolution. These findings support our hypothesis that the SPC-forming protein SPC33 is involved in morphological differences of SPCs and morphological differentiation of perforate SPC is determined by a few key sites. Especially in the site 357, The nucleic acid difference at the 3rd position of Punst (*Punctularia strigosozonata*) Glotr (*Gloeophyllum trabeum*), and Phaca (*Phanerochaete carnosa*) likely occurred after they acquired the CGN codon. This means that parallel evolution occurred in both nucleotide and amino acid sequences of *spc33* during morphological convergence of perforate SPCs. In other words, during the morphological convergence of perforate SPC, convergent evolution at the same site in same gene was observed. Although further validation is needed since we cannot rule out the possibility that other genetic substrates may drove the emergence of Perforate SPCs, here we say the same site-level substitutions can be involved in the morphological convergence of Perforate SPCs. Therefore, the morphological difference between imperforate SPC and perforate SPC may be regulated by a limited number of key sites. If so, there may be few evolutionary paths leading to the change to Perforate SPC.

In conclusion, we successfully found suggestive evidence for the correlation of convergence between amino acid sequence evolution and the morphological evolution of perforate SPCs. Nucleotide and amino acid changes in the same sequence of the same gene corresponded to morphological convergence. This suggests that limited pattern of sequence evolution causes phenotypic convergence. We also showed that present/absent pattern of *spc33* and the short region of *spc33* containing sites with perfect substitution pattern directly linked with morphological evolution of SPC. These findings are a first step for clarifying the genetic basis of SPC morphological evolution. The methods used also made the convergent substitution pattern more easily detectable. Our findings contribute to both clarifying the genetic basis of morphological convergence and demonstrate how genome-wide surveys can be used to elucidate fungal evolutionary morphology.

## Acknowledgements

We would like to thank the people of the Laboratory for DNA Data Analysis at the National Institute of Genetics for their valuable comments on our study. Some computation in this work was performed using the NIG supercomputer at ROIS National Institute of Genetics. We also thank Mallory Eckstut, PhD, from Edanz (https://jp.edanz.com/ac) for editing a draft of this manuscript. TI was supported by JSPS KAKENHI grant number 18J13859 and SOKENDAI future scientist award 2017.

---

**Supplementary Table 1.** AU test result of *spc33*.

| Topology | logL | p-AU |
| --- | --- | --- |
| Species phylogeny | -6631.84 | 0.024 |
| Gene tree | -6624.07 | 0.499 |

**Supplementary Fig. 1.**
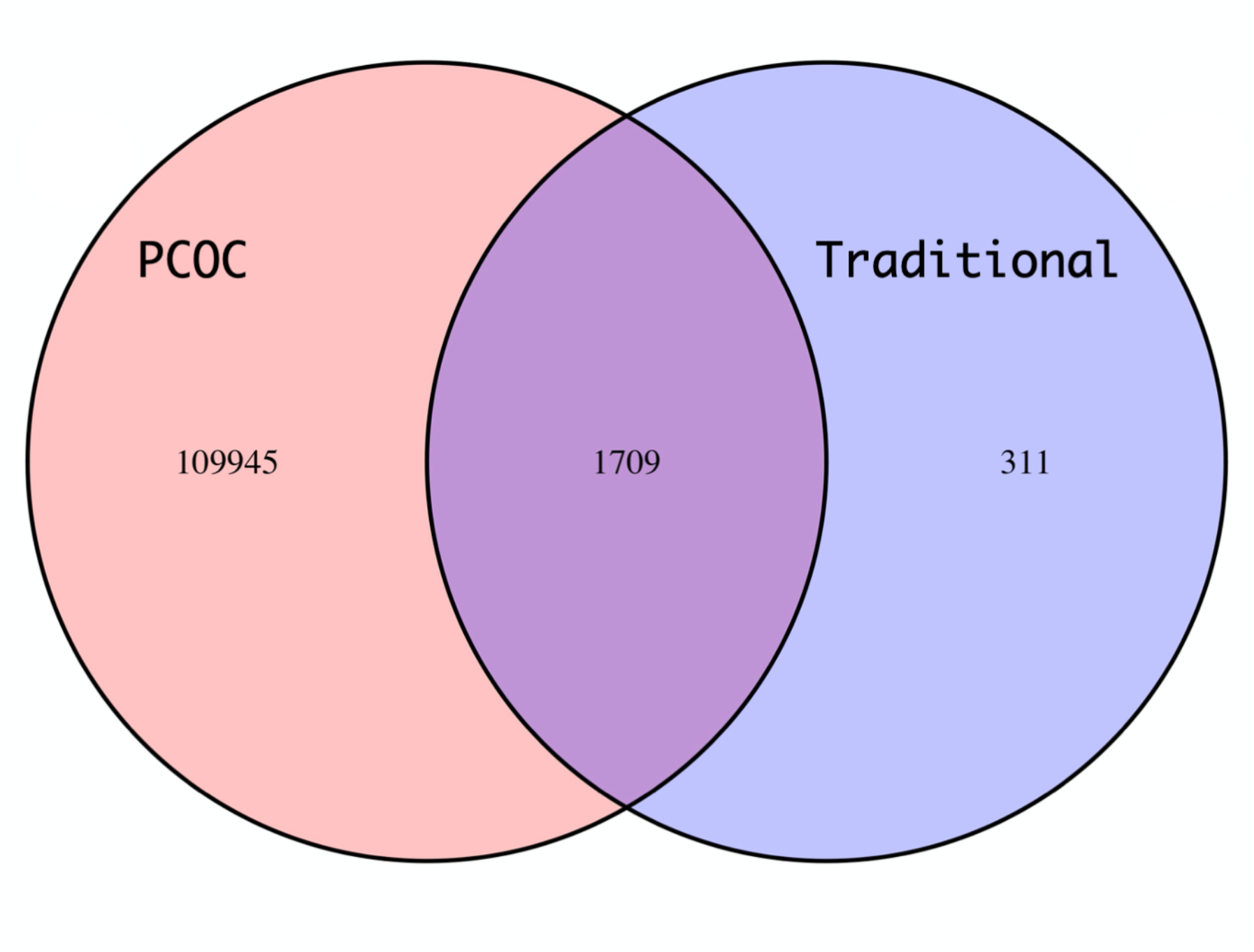
Venn diagram of the number of convergent substitution sites in all orthologous genes detected by our traditional analysis (right) and PCOC analysis (left).

**Supplementary Fig. 2.**
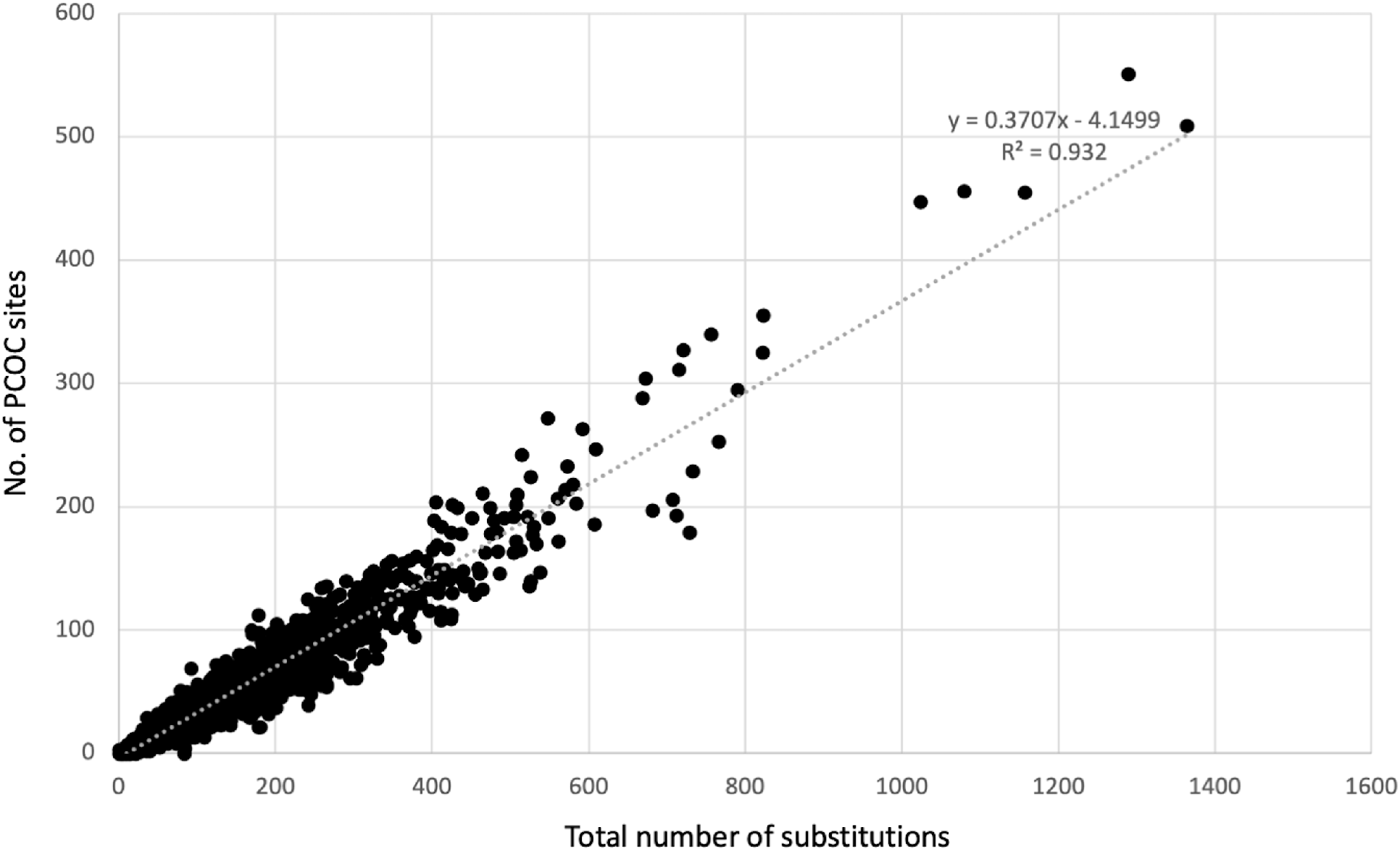
Relationship between the number of convergent sites and the total number of substitutions. The black dots represent the correlation between the number of whole substitutions and PCOC sites for each gene. Gray line indicates approximate straight line of the data.

**Supplementary Fig. 3.**
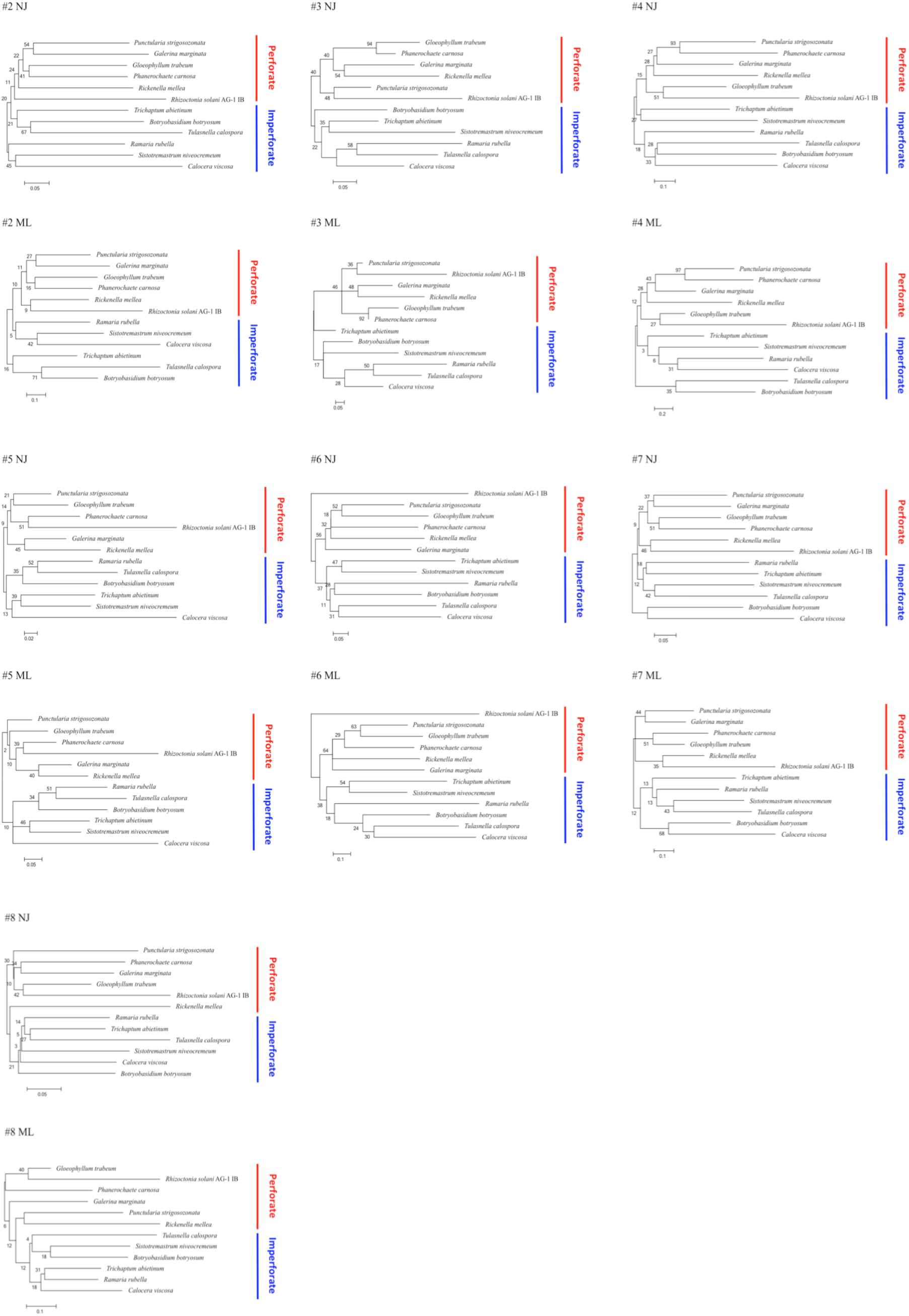
Phylogenetic relationships of correlated genes #2–#8 shown in Table 2. The gene trees were produced using the neighbor-joining (NJ) and maximum-likelihood (ML) methods.

**Supplementary Fig. 4.**
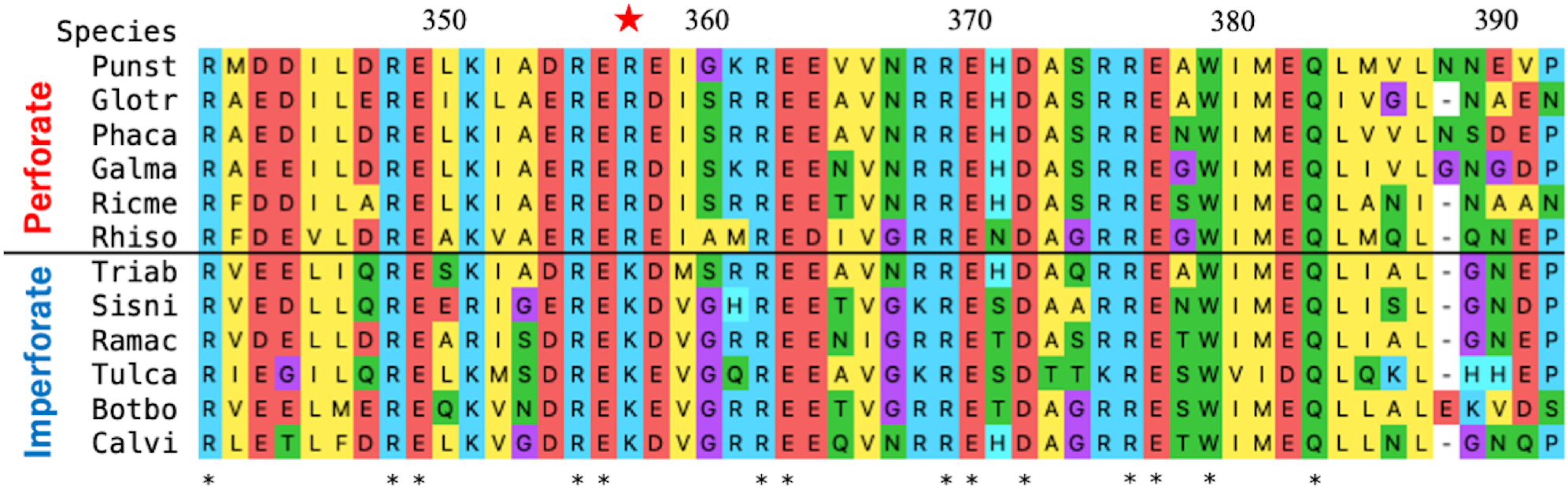
Multiple alignment of site 357 (red star) and neighboring sites in *spc33*.

